# Assessment of whether published non-Cochrane systematic reviews of nursing follow the review protocols registered in the International Prospective Register of Systematic Reviews (PROSPERO): A comparative study

**DOI:** 10.1101/2020.04.14.040865

**Authors:** Kaiyan Hu, Fan Mei, Qianqian Gao, Li Zhao, Fei Chen, Qinghua Liu, Lingling Suo, Yuxia Ma, Ting Zhang, Weiyi Zhang, Bing Zhao, Bin Ma

## Abstract

**Purpose:** We compare published non-Cochrane reviews of nursing with their pre-registered protocols on PROSPERO to quantify the prevalence of differences and the extent to which the differences were explained.

**Methods:** We searched for protocols and their corresponding reviews in PROSPERO’s nursing group that were “completed and published” from inception to September 8^th^, 2019. Two authors independently identified differences and classified the difference as none, partial, or complete, and determined if the existed differences had been explained. Frequency (n), percentage (%), median, and inter-quartile ranges were used to analyze the extent of differences and explanations.

**Results:** We identified 22 pre-registered protocols and their reviews. All 22 pairs (100%) exhibited differences. Eighteen pairs (82%) showed differences in at least six methodological sections, while 21 pairs (95%) involved completed difference in at least one section. The median number of differences per review was 8.00 (upper quartile = 6.00, lower quartile = 9.75). The differences involved all 13 compared methodology-related sections. Only 5 (3%) of all differences were explained in the systematic reviews.

**Conclusions:** We observed widespread differences between non-Cochrane reviews of nursing and their protocols recorded in PROSPERO, with relatively few explanations for the changes. Measures including establishing a new item in the reporting guideline of systematic reviews to guide reporting and explaining the reasons for differences between protocols and systematic reviews or even requiring authors to do so at the Journal’s author guideline are recommended to improve transparency.

## Introduction

Systematic reviews are recognized as being vitally important for evidence-based health care and guide clinical decision-making [1]. However, due to the nature of retrospective design, selective inclusion and reporting of outcomes must be addressed at the level of systematic review [2–4].

To ensure transparency in the assembly and writing of systematic reviews, some organizations, including the Cochrane Collaboration Organization [3] and the Joanna Briggs Institute [5], require registration of the review title and submission of the protocol, which documents the methods and planning of the review. However, these organizations produce only a relatively small proportion of published systematic reviews [1], and mandatory registration for most systematic reviews is lacking. In 2010, PRISMA issued a statement advocating registration of systematic review protocols and this was followed by the establishment of the International Prospective Register of Systematic Reviews (PROSPERO) by the United Kingdom Centre for Reviews and Dissemination in 2011[6]. Their objectives are to effectively fill the vacant position of the systematic review registration and provide a new registration platform for non-Cochrane systematic reviews with which they expect to improve the quality of these reviews [6–7]. For registration on PROSPERO, authors are required to submit information on the design and conduct of the review, including information on 22 mandatory and 18 optional items. Changes, amendments and updates can be made to a published protocol [8].

In some instances, there are valid reasons for protocol alteration, provided the results remain unknown. Legitimate modifications, for example, may extend the scope of searches to include older or newer studies, expand eligibility criteria that were possibly too narrow, or include additional analyses if the primary analysis suggests that this might be warranted. When such alterations are made it is important that they be fully documented and explained to avoid introducing bias into the study [3–4]. Wherever possible, protocol changes should be avoided in order to ensure that research methodology is not changed in response to unexpected results [9]. It is particularly important that these changes be avoided during the data gathering and analysis steps of the study [9–10].

The number of systematic reviews indexed by MEDLINE has tripled over the past decade, with more than 8,000 published in 2016 alone [1]. It is also noticeable that there are differences between the final published reviews and the previously published protocols. Silagy, Middleton, and Hopewell (2002) reported that 43 out of 47 Cochrane systematic reviews contained major methodological changes [11]. Up to September 8th, 2019, there was a total of 851 systematic reviews registered in PROSPERO’s nursing group of which 27 were completed and published [12]. Yet to date, no study has compared the texts of published protocols in PROSPERO’s nursing group with their corresponding published reviews. While the Cochrane library conducts a rigorous assessment of registered protocols, PROSPERO does not and has no peer review or quality assessment [8]. With PROSPERO emerging as one of the major registration platforms for non-Cochrane systematic reviews, it is important to assess whether the protocols registered on PROSPERO have improved the transparency of non-Cochrane systematic reviews, specifically considering reviews in nursing. As such, our primary objective was to investigate and quantify differences in methodology-related sections between non-Cochrane systematic reviews and their pre-registered protocols on PROSPERO’s nursing group. Our secondary objective was to determine the extent to which these changes were reported and explained in the published systematic reviews.

## Methods

### Protocol and review identification

Published systematic reviews of both qualitative and quantitative research (interventions and observations) that had been pre-registered on PROSPERO’s nursing group were identified. This was done by electronic searching for all non-Cochrane nursing protocols on the PROSPERO platform that were “completed and published” from inception to September 8th, 2019. These records usually included citations and links to the final publication; if these were absent or invalid, the open databases (PubMed, Embase and Web of Science) were searched using the review title. The most recent version of the protocol was downloaded after a published review was identified. Two reviewers independently (K.Y.H and L.Z) examined the text of all “pairs” to exclude Cochrane reviews, JBI reviews, and reviews included non-clinical studies. We did not limit the language of published systematic reviews.

### Assessment

For each methodology-related section, two reviewers (K.Y.H. and F.M.) were required to compare the relevant text (including related supplementary files) and assess independently whether there were differences between the systematic reviews and their protocols in all methodology-related sections and, if so, judged and classified the differences as partial or complete. Any divergences in opinion between the two reviewers were resolved by consultation with the third reviewer (B.M.). Initially, a random sample of ten included systematic reviews was simultaneously assessed by the two reviewers (K.Y.H. and F.M.), with the formal assessment only commencing when significant (>90%) agreement was reached. Differences in any of the following 13 methodology-related sections were assessed: Review question, Search strategy, Participant(s)/population, Intervention(s)/exposure(s), Comparator(s)/control, Type of study design, Main outcome(s), Additional outcome(s), Study selection, Data extraction, Risk of bias assessment, Data synthesis, and Subgroups analysis. These 13 sections are the mandatory registration entries required by PROSPERO.

A language expert (L.L.S.) was employed to translate the non-English reviews. The translator has a multilingual background and is a professional medical translator. Before translation, we introduced the purpose and method of the research to the translator, and at the same time trained her with the systematic reviews production process. Two reviewers (K.Y.H. and Q.Q.G.) and the language expert repeatedly proofread the texts in the process of translation to ensure accuracy of the translation.

When differences between a systematic review and its protocol were observed, two authors (K.Y.H. and F.M.) independently investigated whether there were amendments to the protocol or acknowledgment and explanation of the alterations in the published review. Consensus between both authors was required on whether differences were adequately explained in the publication.

### Classification of the extent of difference

Differences were judged and classified using an internal guideline that was developed, independently pilot-tested (in n=10 pairs), and revised by two authors (K.Y.H. and B.M.) (S1 Appendix). The categories used were: no difference (when the systematic review and protocol matched or when a section was either not applicable or not reported), partial difference (when a section was either not fully specified in the systematic review or the protocol, or when minor differences were observed between parts of sections), and complete difference (when aspects of a section were modified in the systematic review or when a section was omitted from either the protocol or the review). If judgments differed on a section, we chose the ‘‘worst case’’ (e.g., when we observed a partial and a complete difference, we classified the compared section as having a complete difference). If there were more than two partial differences concerning the one section, we also classified it as having a complete difference.

### Statistical methods

Statistical analysis was performed with Microsoft Excel 2016 software. Frequency (n), percentage (%), median and Inter-quartile range were used to analyze the extent of difference in each compared methodology-related section of the reviews and the extent of reporting and explanation for the differences in the systematic reviews.

## Results

### Included studies

From inception of the platform to September 8th, 2019, 851 non-Cochrane protocols were registered on PROSPERO’s nursing group, with 27 reviews completed and published. Of these 27 reviews, two were excluded because they were JBI reviews, two were excluded because they were assessments of health technology, and one was excluded because it was a clinical practice guideline. Twenty-two systematic reviews were eligible and included (Figure 1. Flow diagram for the identification and selection of eligible systematic reviews in this study.); two of these were in German, one in Farsi, and 19 in English. All excluded and included reviews were listed in S2 Appendix.

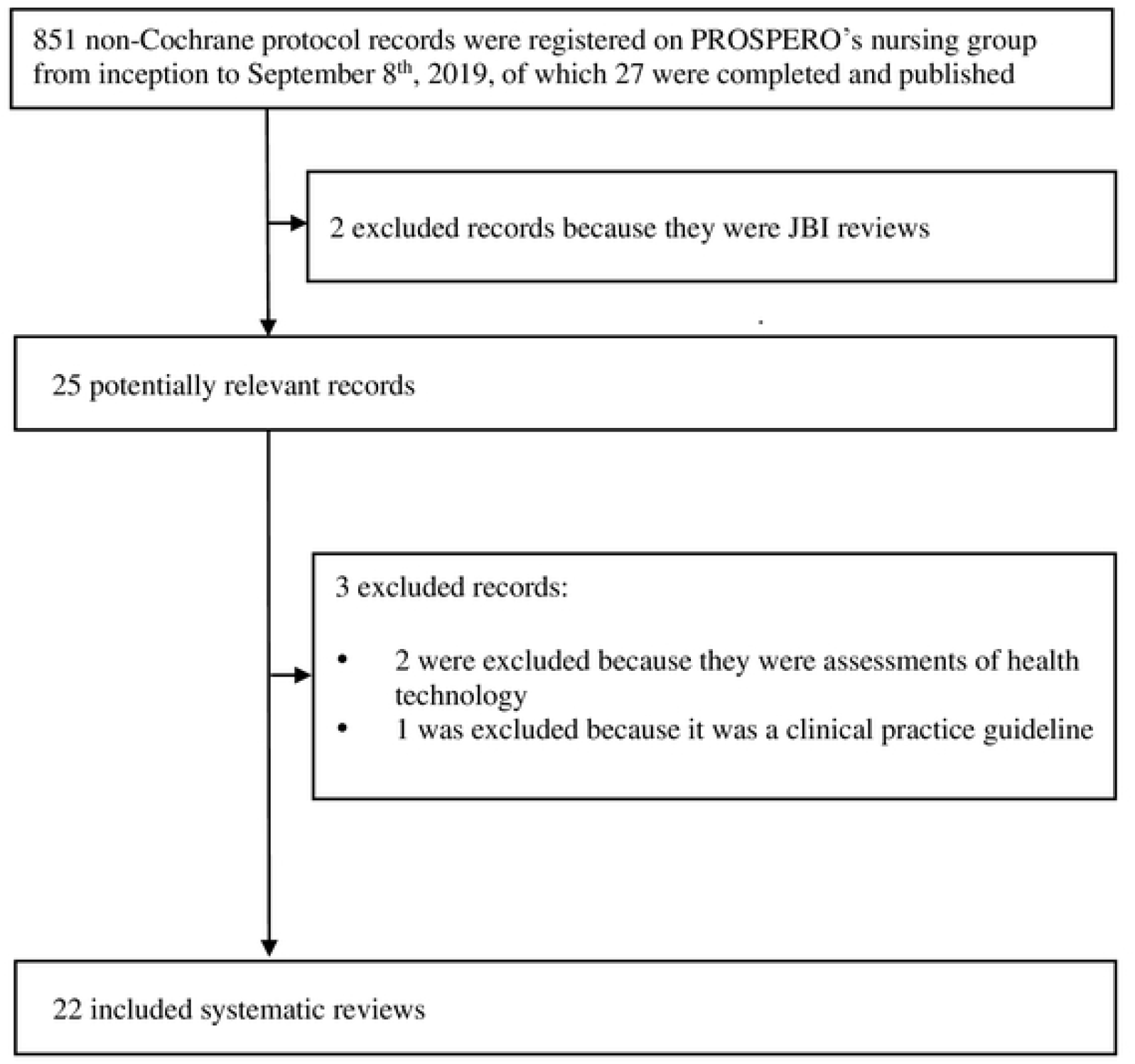

### Results of comparisons

All differences between the systematic reviews and their protocols are listed in S3 Appendix. These differences were seen in all the included systematic reviews and in all 13 of the methodology-related sections. We observed that many protocols were deficient in key information such as details of participants, intervention/exposure, comparator, and outcomes. More alarmingly, key information was often missing in mandatory fields such as “Search strategy”, “Study selection”, “Data extraction”, “Risk of bias assessment”, and “Data synthesis”.

### Extent of changes

All 22 pairs (100%) exhibited differences, and the extent pf changes was categorized based on data in S3 Appendix. At least six sections involved differences (completed or/and partial) in 18 pairs (82%). At least four sections occurred partial difference in 13 pairs (59%), and at least one section occurred complete difference in 21 pairs (95%). The median number of difference peer review was 8.00 (upper quartile = 6.00, lower quartile = 9.75). The prevalence of difference in each pair was shown in Fig. 1. (Figure 2. The prevalence of difference in each pair)

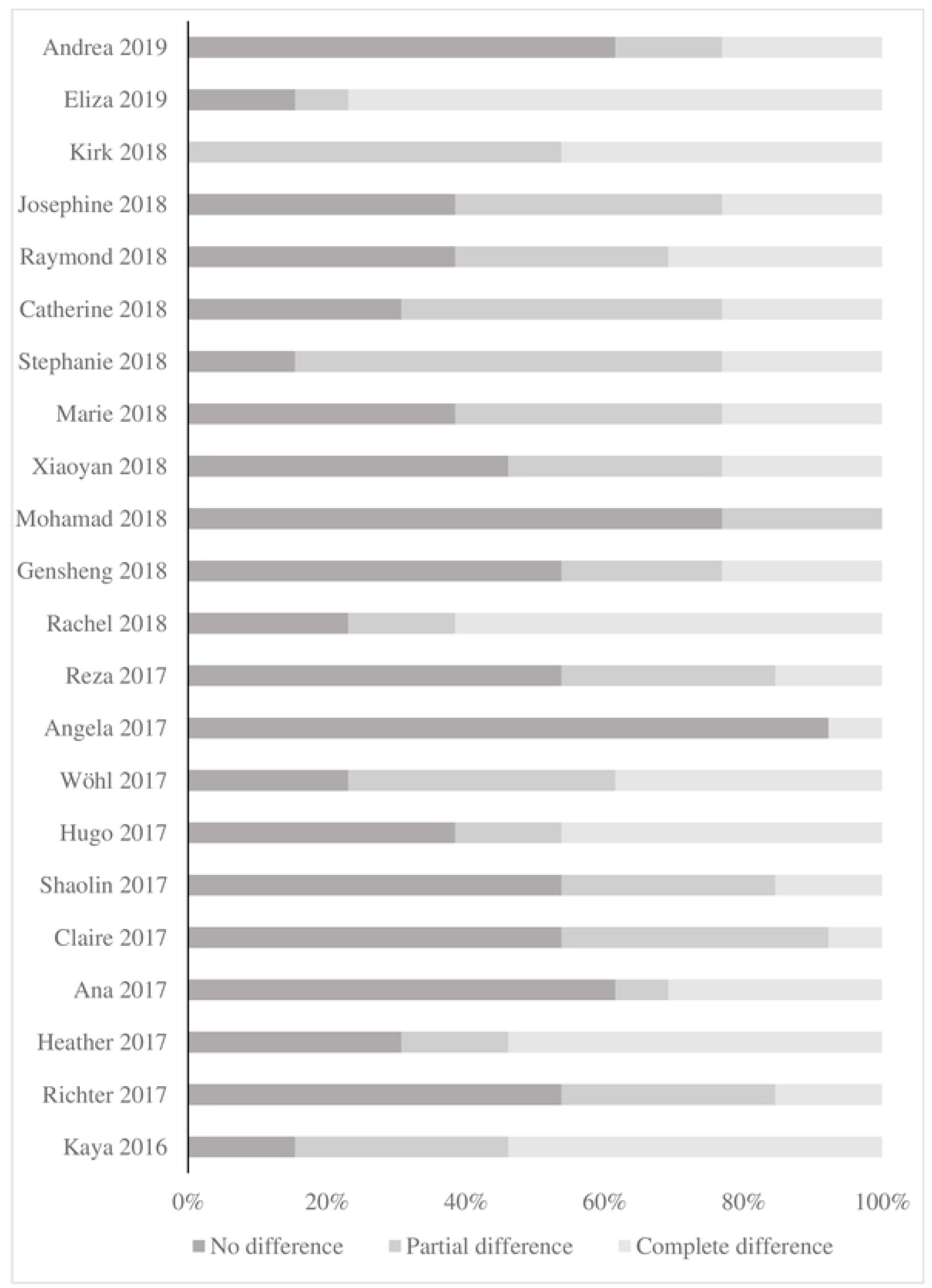

Differences involved all 13 methodology-related sections. All pairs (100%) showed differences (completed or/and partial) in “Search strategy”. Half of the pairs differed (completed or/and partial) in the sections “Type of study design”, “Intervention(s)/exposure(s)”, “Participant(s)/population”, “Main outcome(s)”, “Search strategy”, “Study selection”, “Data extraction”, “Risk of bias assessment”, and “Data synthesis”. Furthermore, there were partial differences in half of the pairs regarding the sections “Data synthesis”, “ Risk of bias assessment”, and “Intervention(s)/exposure(s)”; there were complete differences in half of the pairs regarding the sections “Search strategy” and “Data extraction”. The prevalence of difference in each section was shown in Fig. 2.(Figure 3. The prevalence of difference in each compared methodology-related section)

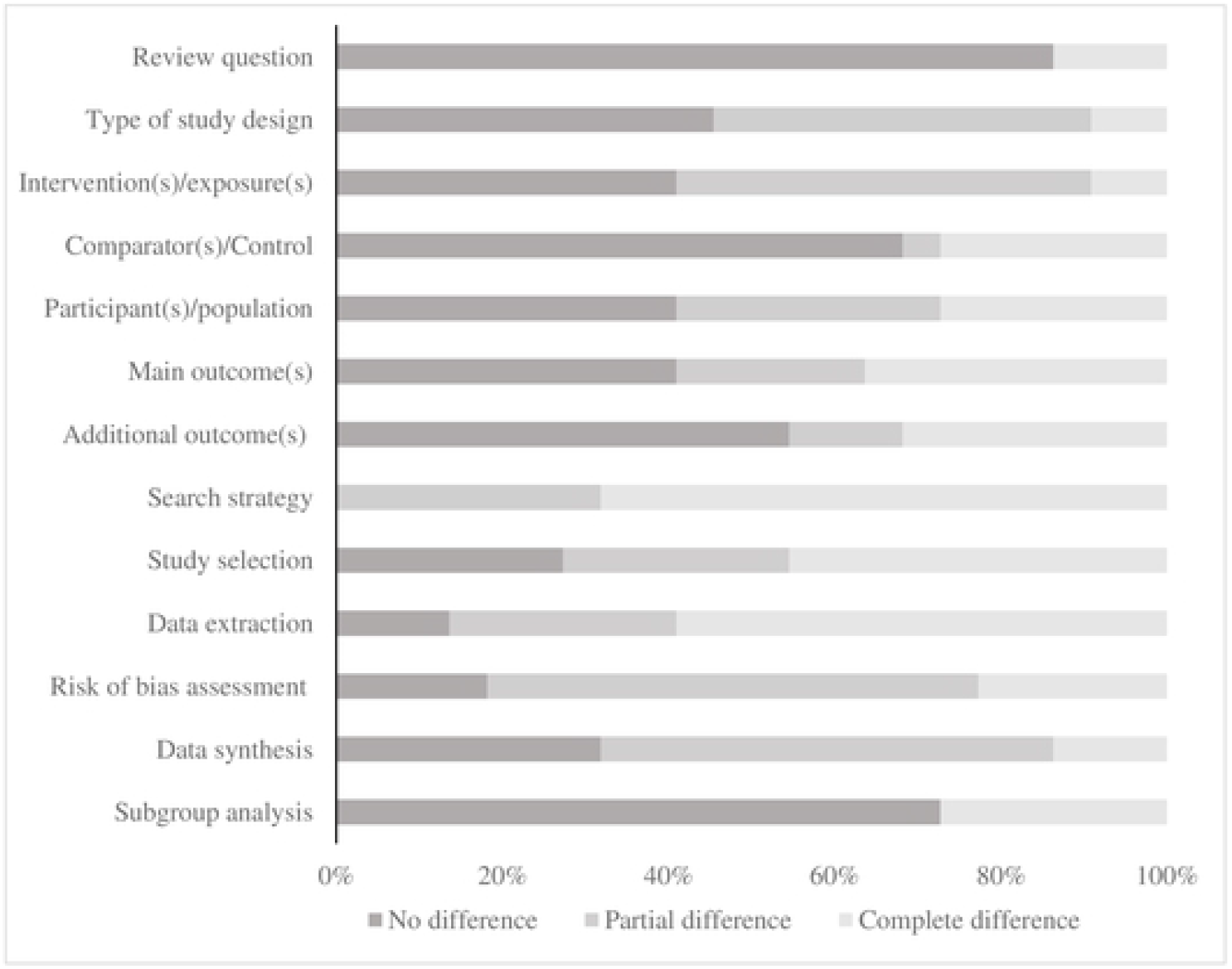

### Reporting and explanation of differences

Of all 286 judgments (22 pairs times 13 sections), 167 involved differences. Of those, 81 were judged to be partial differences and 86 to be complete differences. Only 5 (3%) of all differences were explained in the systematic review, and these only occurred in “Data synthesis” and “Subgroup analysis”. The prevalence of explanation for all differences was shown in Fig. 3. (Figure 4. The extent of explanation for all differences)

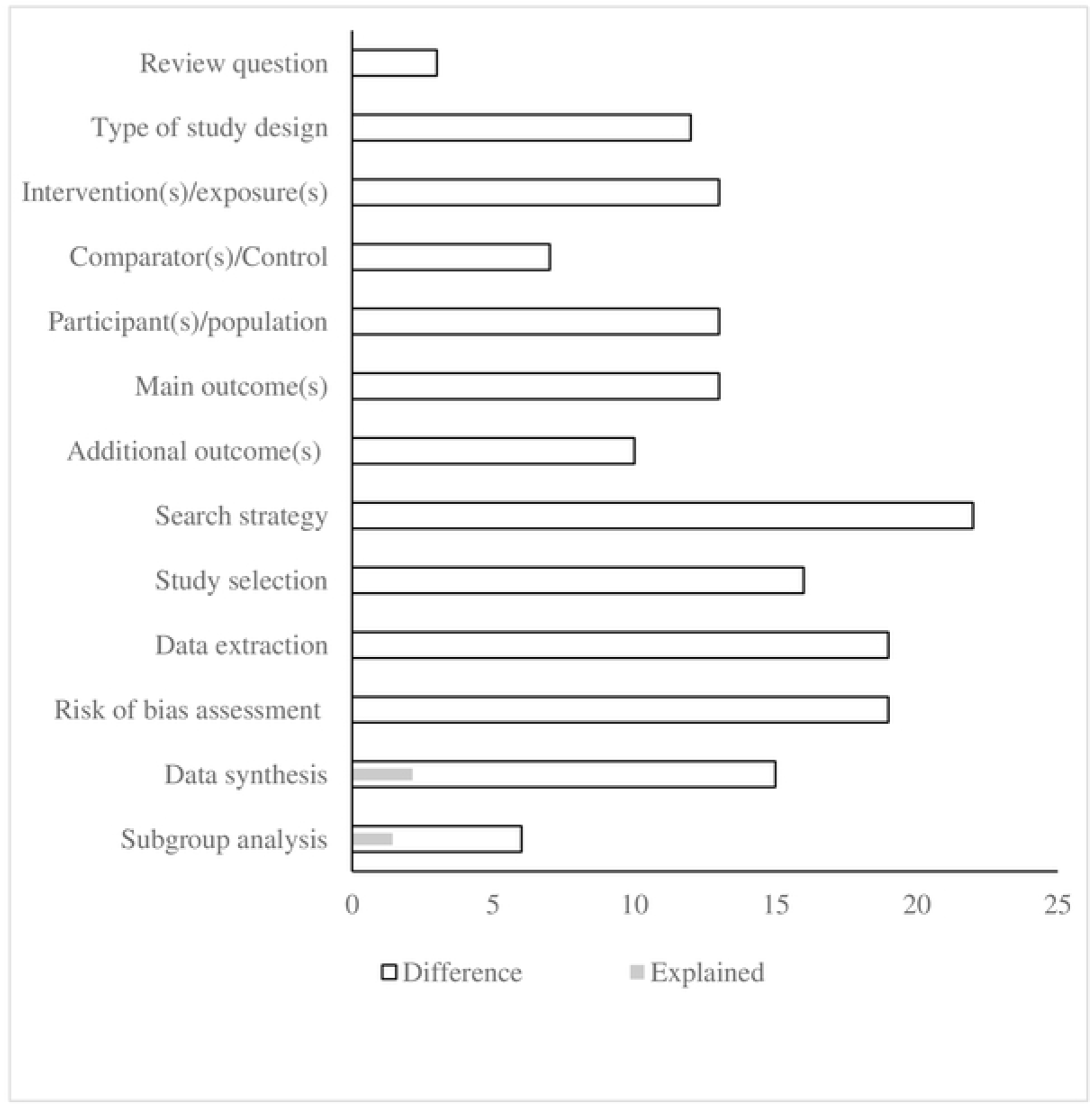

## Discussion

This is a descriptive investigation of all non-Cochrane review protocols registered on PROSPERO’s nursing group and a comparison with their corresponding published full-text reviews. We found alterations in all pairs and showed that these alterations involved all 13 compared methodology-related sections. However, only 3% of all alterations were explained in the systematic reviews.

### The impact of changes

While some alterations may clearly impact the methodological and reporting quality, as well as introduce risk of bias, in the published systematic review, this may not necessarily be apparent in all cases [13]. An example is the application of limits to language of publication and databases, which might result in a high risk of bias for a particular area. Modification of participants, interventions, comparison, outcomes, study design, and subgroup analysis are not as obvious in terms of potential sources of bias or manipulation. Ideally, the review should be conducted in strict accordance with the published protocol, as this would reduce the chances of bias resulting from possible conscious or unconscious manipulation to reach an anticipated conclusion [9]. Nevertheless, in some cases, there may be valid reasons for modifying protocols. Such legitimate modifications may include broadening the search period to include newer or older studies, expanding eligibility criteria that may have been found too narrow, or the addition of further analyses if the primary analyses suggest that these are warranted. However, it is necessary to describe the alterations and explain the rationale for including them in the review to reinforce the transparency of the systematic review process [3–4].

We found that all pairs (100%) underwent some alterations during the research process with only 3% of these differences explained in the systematic reviews. Thus, the transparency of non-Cochrane systematic reviews of nursing registered on PROSPERO’s nursing group is deemed inadequate. A recent study compared 80 non-Cochrane systematic reviews with their published protocols, and found that almost all (92.5%) differed from their protocols in at least one of the methods-related “Preferred reporting items for systematic review and meta-analysis protocols” (PRISMA-P) items and their subcategories, with only 7% of these providing an explanation [14]. Similar discrepancies between non-Cochrane systematic reviews and their PROSPERO protocols have been reported [15–18]. Although these discrepancies appear to be fairly common, reasons for the differences are rarely addressed.

### Improving the transparency

The responsibility of authors to follow their protocols or to clearly describe and explain the reasons for any alterations should be highlighted. Authors of systematic reviews should be encouraged to record changes to protocols, with explanations, on PROSPERO. We also suggest adding a new item to “Preferred Reporting Items for Systematic reviews and Meta-Analyses”(PRISMA), “Meta-analysis of Observational Studies in Epidemiology” (MOOSE), and “Enhancing transparency in reporting the synthesis of qualitative research”(ENTREQ) to guide authors reporting the alterations (with explanations) between protocols and systematic reviews. This would also allow readers and users of systematic reviews to be aware of the implications of reviews not adhering to the protocols. In addition, journals could include a request to authors in their author guidelines to provide details of any changes (with appropriate reasons) made to the original protocol in an appendix to the published review. This would allow editors or peer reviewers to compare manuscripts systematic reviews with their protocols and check discrepancies with the authors. Regulation at a higher level, such as the International Medical Journal Editors’ Committee (ICJE), is also recommended to improve transparency and uniformity in biomedical publishing.

### The strengths and limitations of this study

This is the first study to compare non-Cochrane systematic reviews with their protocols registered on PROSPERO’s nursing group regarding differences in all methodology-related sections, as opposed to only differences in predefined outcomes. We acknowledge several limitations to our study. Firstly, we identified the published systematic reviews by using the final publication detail of PROSPERO’s record. It is therefore possible that the reviews could be misses if the information was not updated. Additionally, although we did not restrict the number of reviews based on language or date of protocol registration, the sample size was still relatively small. Secondly, we did not verify the differences with the authors due to limited resources. Our study was based solely on the reporting of the systematic reviews and their protocols, resulting in decisions on differences being affected by the quality of the reporting. Implication for future research A necessary focus for future research is the reporting quality of the protocol registered on PROSPERO’s nursing group. The protocol allows comparison, with complete reviews to determine possible reporting bias [13]. The reporting quality thus affects conclusions on differences between the protocol and the review, influencing the transparency of the review.

Currently, there is no peer review or quality control of PROSPERO’s protocols, increasing the probability of poor reporting quality [8]. It is also necessary to verify differences and explanations with authors in order to have an understanding of the reasons for the changes and their impact on the quality of the final review. The question of whether altering a specific aspect of a methodology-related section is associated with higher or lower methodological quality should also be investigated.

## Conclusions

Differences between the non-Cochrane systematic reviews of nursing and their protocols recorded on PROSPERO’s nursing group were widespread, with relatively few explanations. We recommend measures to improve transparency of non-Cochrane systematic reviews of nursing, specifically, establishing a new item in PRISMA, MOOSE, and ENTREQ to guide reporting and explaining the reasons for differences between protocols and systematic reviews or even requiring authors to do so at the Journal’s author guideline.

## Acknowledgements

This research did not receive any specific grant from funding agencies in the public, commercial, or not-for-profit sectors.

## Supporting information captions

S1 Appendix. An internal guideline.(docx)

S2 Appendix. All included and excluded reviews. (docx)

S3 Appendix. All differences between the systematic reviews and their protocols. (xlsx)

